# High density probes reveal medullary seizure and rapid medullary shutdown in a model of fatal apnea in seizure

**DOI:** 10.1101/2025.02.10.637524

**Authors:** Ryan Budde, Laura RoaFiore, Pedro Irazoqui

## Abstract

**Objective:** Sudden unexpected death in epilepsy (SUDEP) is suggested to be a cardiorespiratory collapse that occurs shortly after a seizure. Prior work in rats suggests that reflexive apneas (produced by stimulation of trigeminal or vagal peripheral sensory targets) is highly fatal during seizure but well tolerated otherwise. These reflexes share network connectivity in the medulla, particularly the caudal solitary nucleus (NTS) and ventral respiratory column (VRC), and possibly other intermediate structures. We sought to observe the electrographic activity in these regions.

**Methods:** We use urethane anesthetized long evans rats. We utilized either 125 *μ*m silver wire in the caudal NTS or a Neuropixel 1.0 probe along a dorsoventral trajectory that spanned the caudal NTS to the VRC. We additionally recorded cardiorespiratory activity via several methods. We induced a reflexive apnea – the diving reflex – by nasal irrigation of cold water for several seconds, which produces a period of apnea, then gasping, and then a gradual return to eupnea. We repeated several trials while the animal was healthy and subsequently induced continuous seizure activity with kainate and repeated the reflexes, which are ultimately fatal during seizure.

**Results:** Seizure activity confounds many established methods of analyzing high-density single unit data such as provided by Neuropixels probes, and so our analyses focus on averaging responses over larger anatomical regions (120 *μ*m) covering small populations of neurons. Seizure produces broad increases in neuronal activity across the medullary tract, which by itself is not dangerous. Ictal reflexive apneas were broadly more inhibitory (producing a reduction in firing rate) than they were preictally, and fatal ictal responses resulted in a very rapid shutdown of all medullary activity. We only rarely observed ictal central apneas (apneas with no apparent stimuli), but when we did they were apparently safe, always survived, and produced no significant change in network activity (neither increase nor decrease).

**Conclusions:** These data support the theory that central apnea events in seizure are relatively safe as we observed they produce little change in the medullary tract network, while stimuli-induced-reflexive-apneas are dangerous because they produce profound quieting across respiratory centers. Our data suggest that seizure spreads to this medullary tract at approximately the same rate and intensity as forebrain, as previously described in this model. These data are supportive of SUDEP mechanisms involving brainstem inhibition as a primary cause, such as spreading depolarization waves. These findings likely extend beyond nasal irrigation to any sensory reflexive apnea caused by airway irritation of any kind, and may bear relevance to similar deaths seen in infants.

## 1 Introduction

Sudden unexpected death in epilepsy (SUDEP) is a fatal complication of epilepsy, and is predicted to be a respiratory collapse that occurs shortly after a seizure (Ryvlin et al. 2013; Verducci et al. 2019). In SUDEP the known risk factors lack predictive power (Thurman David, Hesdorffer Dale, and French Jacqueline 2014), but recent work has suggested that there may be measurable changes in the patient in the weeks-to-months leading up to the fatal event (Simeone et al. 2024), including respiratory impairments seen even outside of seizure (Sainju, Dragon, Winnike, Vilella, et al. 2023; Sainju, Dragon, Winnike, Nashelsky, et al. 2019). Additional work has suggested several mechanisms by which seizure may produce widespread respiratory inhibition, either as a fundamental mechanism of hyperactivity (Aiba and Noebels 2015), or from seizure spread from the forebrain down to medullary respiratory centers (Nobis et al. 2019; Faingold and Feng 2023). We are interested in the intersection of these two theories – acquired impairments to respiratory circuits that may increase the likelihood of a rare fatal response. Our work has focused on reflexive apnea circuits contained entirely in the hindbrain and likely confined to just the caudal medulla. We were initially intrigued by the observations of laryngospasm in the acute rat kainate model (Nakase et al. 2016). Over several studies we established the following: we observed that, in the kainate model, seizure causes stomach acid overproduction and air swallowing; these factors cause stomach acid to reflux up the esophagus and into the airway if the animal is prone; vagal laryngeal chemoreceptors respond strongly to the acid, triggering laryngospasm (an obstructive apnea during which there is movement of the diaphragm); laryngospasm is brief and then animals display a strong central apnea (no movement of diaphragm) that is highly fatal (R. B. Budde, Arafat, et al. 2018; R. B. Budde, Pederson, et al. 2020; Mandal et al. 2021). Critically, this irritant-induced-reflexive-apnea is only fatal in seizure; nonseizing animals, if subjected to artificial acid reflux in the airway, initially display a similar apnea response but always start breathing again and survive. Seizing animals display low respiratory drive and die. These observations led us to a wealth of literature of apnea control in sudden infant death syndrome (SIDS), particularly laryngeal chemosensation in both lambs and humans which could produce profound apneas in infants but not older subjects or adults (Johnson, Dawes, and Robinson 1972; McGraw 1939; Goksör, Rosengren, and Wennergren 2002; Marchal, Corke, and Sundell 1982; Davies, Koenig, and Thach 1988; Donnelly, Bartlett, and Leiter 2016; Scadding et al. 2014). Some of these papers also point toward dangerous apneas in infant swimming, which can trigger a diving reflex. Interestingly, the diving reflex apneas, suggested to be sometimes dangerous in infants, feature trigeminal stimulation of the face or nasal epithelium, not vagal laryngeal airway sensation (Pedroso et al. 2012; Panneton and Gan 2020). We next tested this trigeminal stimulus in our acute kainate experiments and found that it, too, was highly fatal in seizure but never in controls (Biggs, R. Budde, et al. 2021). Furthermore, the responses to the different stimuli (vagal vs. trigeminal) were largely indistinguishable from one another, and so we hypothesized they shared a downstream apnea network. Additional literature in diving, breath-holding, and respiratory sudden deaths implied carotid body function (Lugliani et al. 1971; Powell, Milsom, and Mitchell 1998), and, indeed, the glossopharyngeal carotid bodies and the vagal laryngeal chemoreceptors share terminal projections in the caudal NTS (Mifflin 1996). The diving reflex apnea was previously never fatal in our healthy controls; however, this reflex acquired moderate mortality with unilateral carotid body denervation and extreme mortality with bilateral denervation, all *without* seizure (Biggs, R. B. Budde, et al. 2022). The deaths from reflexive apnea with denervation (i.e., pathway inhibition) and no seizure were essentially identical to those with intact nerves and seizure. Furthermore, in this study we demonstrated that electrical stimulation (i.e., pathway excitation) reduced mortality in apnea trials during seizure. Therefore in these fatal apneas the effects of seizure were not distinguishable from carotid body inhibition.

This prior work demonstrates that three separate apneic pathways (trigeminal, vagal, and glossopharyngeal) display extremely conserved pathophysiology in our model of sudden respiratory deaths in seizure. These three pathways also display literature of plasticity and gradually acquired impairments, possibly linking the recent human observations of acquired respiratory impairment. We therefore hypothesize that these pathways share a significant downstream apnea network which can be fatal, especially in seizure, and this downstream network may contribute to the sudden death mechanism. In this study we record from various medullary regions possibly associated with these pathways. Prior work has demonstrated that the carotid body hypoxia sensor and laryngeal chemoreceptors project to the caudal solitary nucleus (NTS) (Mifflin 1996) and the nasal epithelium trigeminal receptors project to the caudal spinal trigeminal (Panneton and Gan 2020). We additionally know that these circuits must ultimately modify the ventral respiratory column (VRC) – a group of critical respiratory structures in the ventrolateral medulla – and particularly the preBötzinger complex which generates each inspiratory effort. Current literature does not establish which other regions may act as intermediaries or if these inputs project immediately to the VRC from the primary input. Therefore, in this study we utilize some of our previous methods (urethane-anesthetized rats, with kainate-induced seizures, and apneas induced with the trigeminal reflex via nasal irrigation with water) with additional recording of a medullary tract. We primarily utilize Neuropixels probes, multi-electrode array probes that allow us to record activity along a 3.8 mm dorsoventral tract, allowing us to capture activity along a section of the NTS, some intermediate structures, and the VRC. Our methods cannot validate the recording of exact structures, especially as some regions like the preBötzinger complex lack specific markers. Therefore we report on more general patterns of activity.

These experiments also allow us to probe critical SUDEP questions – how significantly does seizure spread to the medulla, and how do medullary respiratory centers behave during respiratory deaths? There is a lack of literature of how often seizure spreads to the medulla, and our anecdotal estimates from clinical colleagues range from “almost never” to “all the time.” Further, if seizure spreads to the medulla, how dangerous is it? Do fatal apneas onset very shortly after medullary seizure, or can the medulla endure seizure for an extended period of time without danger? Additionally, there is little literature on the behaviors of medullary structures in ictal deaths. One might assume a respiratory death would mimic coma or chronically depressed consciousness (vegetative state), in which the brainstem perseveres and remains the last active brain region. However some animal data suggests the brainstem may in fact be the source of the SUDEP brain shutdown (Aiba and Noebels 2015). Lastly, how do different apnea types impact the medulla? Central apneas are common in patients and animals and are overwhelmingly safe. However in the rat kainate model obstructive or reflexive apneas are likely to be fatal. Do different apneas produce different responses in the medulla? And finally, prior data shows the obstructive apneas are only fatal in seizure. Does the response to obstructive apnea change with seizure?

In summary, we find:

1. Seizure activity spreads to the medulla at approximately the same rate and intensity as it does to cortex in this model. Medullary seizure is safe so long as animals do not experience reflexive apneas.
2. During seizure reflexive apneas always shift to become more inhibitory (producing a reduction in firing rate) than they were before seizure.
3. The entire medullary tract shuts down very rapidly in fatal apnea trials, possibly faster than cortex.
4. We often observe swallowing during ictal apnea but not during preictal apnea. Some, but not all, medullary data suggests swallows and gasps may act counter to one another.
5. A small sample size suggests central apneas do not produce dramatic changes in medullary firing.

## 2 Methods

### 2.1 Summary

We performed nonsurvival experiments on young adult rats that were anesthetized and otherwise healthy. We recorded various cardiorespiratory metrics and medullary activity. We apply a diving reflex stimulus by spraying cold water in the nose for a few seconds which then runs out the mouth (it is not aspirated). This stimulus produces a period of apnea and then gasping as the animal recovers over a few minutes. We repeat the stimulus a few times to gather control data, then induce continuous seizure activity, and then repeat the stimulus again in the ictal state.

### 2.2 General procedures

We include data from 17 animals with an additional 29 used in the development of the methods. All surgical procedures were approved by the Johns Hopkins University Animal Care and Use Committee and comply with ARRIVE guidelines. Animals were housed in same sex pairs under a 12:12 light-dark cycle, with ad libitum access to food and water. We include female (*N =* 11, 272 ± 31 g; range 225–353 g; Envigo) and male (*N =* 6, 361 ± 54 g; range 308–455 g; Envigo) animals. We did not observe any significant differences in data conclusions between sexes, and so all data are presented with both sexes grouped together.

We anesthetized animals with urethane (1.5g/kg, i.p.), and used a thermostatically-controlled heating pad to maintain core body temperature near 37°C throughout the experiment. Rats were placed into a stereotaxic frame and headfixed using ear bars, an incisor bar, and a snout clamp with very mild pressure. For nasal irrigation we placed a small silicone cannula in the right nostril to a depth of approximately 4mm with marks to indicate if the tube moved during the experiment. We additionally used a small amount of cyanoacrylate glue around the seal between the cannula and the surrounding mucosa. A 1 inch steel plate was placed on the animal’s skin over the dorsal pelvis using a conductive gel to serve as ground. We next set up the recording of various cardiorespiratory and medullary data as described below. After instrumentation we collected at least 15 minutes of baseline data in each animal.

Each animal was subjected to multiple reflexive apnea trials, both before and during seizure. Before seizure we performed 3–6 trials of approximately 0.8 mL of water over 8 seconds (the 0.1mL/second flow rate is consistent in all cases in all animals). Trial numbers and durations are variable as the animal response is variable and we prioritize at least 3 apneas of at least 3 seconds duration. Immediately after we cease the stimulus we apply mild suction to remove any additional water in the cannula or sinus. The stimulus causes an apnea and gasping recovery, and we wait 5 minutes or until the animal returns to baseline respiratory function before proceeding, whichever is longer. After sufficient preictal data we induce seizure with kainate (10 mg/kg i.p.) and wait 45 – 90 minutes for seizure to develop before proceeding. The time to seizure is assessed primarily by the animal’s respiratory pattern (which acquires a stereotypical pattern of 6–12 rapid shallow breaths punctuated with sighs), the mild onset of tonus across the dorsal musculature (slightly modifying body posture and tone), exophthalmos, and mild clonus of the vibrissae. We then proceed with ictal trials. In each ictal trial we would deliver 0.1 – 2 mL of water via the nasal cannula over 1 – 20 seconds. We varied the duration and volume of water in order to fine tune the apnea intensity. Animals display different sensitivity to the stimulus during seizure, (anecdotally) not correlated to their sensitivity before seizure, and if the stimulus is too strong then the animals may die before we gather sufficient data to draw useful conclusions (2/13 animals died on the first ictal trial even with the mildest stimulus, and therefore not all data comparisons are possible in these animals). Therefore we started with a shorter stimulus and gradually increased the duration. We prioritized the shortest stimulus which still produced apnea for at least 3 seconds after the stimulus is removed. If the first, very mild diving reflex stimulus did not produce an apnea of at least 3 seconds then that trial was not included in data analysis. The preictal duration (8 seconds) was chosen to be an approximately median response of apnea duration over the 1 – 20 second range for comparable data. Eventually one of the ictal apnea trials is fatal, occurring on trial 1–12 in this dataset.

### 2.3 Measures

#### 2.3.1 Medullary

Prep and coordinates: We resected the scalp skin and subcutaneous fascia to reveal the whole dorsal area of the skull. We removed the periosteum and leveled the skull in the horizontal plane between bregma and lambda. We used a dental drill to drill a burr hole approximately 5 mm caudal to the primary window. Next, we used a stainless steel bone screw with soldered pins at the burr hole to serve as a reference to the electrode(s). We then used the dental drill to open the primary cranial window at ML *−*0.50mm, AP *−*12.5mm from bregma. We made a small durotomy using a #11 scalpel in order to facilitate probe insertion, making sure to not disturb brain tissue. The exact implant location was modified to avoid penetrating surface vasculature, which varied in each animal. Both electrode types were inserted at an angle of 10°away from midline. Our prior work has found this trajectory greatly reduces bleeding when targeting medullary structures (Arafat et al. 2019; Jefferys et al. 2019). All probe insertions were at animal left, which was limited by our available stereotaxic and electrode insertion tools. *Neuropixel probe:* We utilized the Neuropixel probe in 13/17 experiments. In order to minimize motion artifact in these animals we added additional skull support beyond the ear bar stereotaxic frame. We used UV cure dental cement (NX3 Nexus) to affix a custom titanium headplate (SendCutSend) to the rostral end of the skull, and used a supporting clamp assembly to more firmly affix the head. We utilized a DiI-dipped Neuropixel 1.0 probe (IMEC) in a motorized micromanipulator (uMP-4, Sensapex). We moved the probe to the center of the cranial window and reference all measurements from the surface of the brain at the beginning of insertion. We first rapidly inserted the probe to a depth of 5400 *μ*m at a rate of 8 – 10 *μ*m per second, and then allowed the probe to settle for approximately 15 minutes. This first insertion did not cross any tissue included in recording or analysis (all cerebellar). Following this wait period we next slowly inserted the probe to its final depth of 9000 – 9150 *μ*m at a rate of 3 *μ*m per second. The final probe depth was chosen to optimize activity near the top and bottom of the probe and was sometimes restricted by skull surface anatomy. We next advanced the probe approximately 100 *μ*m further in depth than the final position, then retracted, to allow for decompression to minimize subsequent tissue movement and relaxation. At the final depth we let the tissue settle for at least 30 minutes before we began baseline recording. We used saline to hydrate the cranial window throughout the experiment. *Single wire:* We used an unsharpened 125 *μ*m silver wire electrode coated with 10 *μ*m parylene insulation. We used these electrodes in 4/17 experiments. We advanced this electrode to approximately 5800 *μ*m and waited at least 15 minutes for the tissue to settle before starting recordings. The amplifiers we used with these electrodes were very sensitive to noise, and so we only recorded medullary activity, respiratory effort assessed via video, and nasal airflow via thermocouple.

#### 2.3.2 Cardiorespiratory

ECG: We used two 25G stainless steel hooked needle electrodes to measure electrocardiography (ECG), placed percutaneously, just caudal to the junction of the upper limb and thorax (in Lead I). Diaphragm EMG: We used two 30G stainless steel needle electrodes to measure electromyography (EMG) of the diaphragm. We placed needles just at or below the most caudal ribs, midline of dorsoventral axis, angled medial and rostral. Pharyngeal EMG: We used two 30G stainless steel needle electrodes in various locations to measure movements of the jaw and associated swallowing behaviors. In some experiments (5/13) we placed electrodes into the geniohyoid and stylohyoid musculature in order to capture deglutition specifically (a true measure of swallowing and movement of the hyoid bone). However this electrode placement also tended to capture activity with respiration, not just swallows. When we observe animals swallowing in experiments there is always visible motion of the lower jaw when a swallow occurs (as confirmed by hyoid muscle recordings) and so in other experiments (8/13) we instead placed the electrodes near the masseter which captured these events without also capturing respiratory activity. Thermocouple: We placed a thermocouple just outside the left nostril in order to measure the air temperature changes associated with respiration. Diaphragm pressure sensor: We placed a small silicone balloon under the diaphragm in the region of greatest displacement with eupnea. We inflated the balloon with air and recorded the pressure inside to measure respiratory displacement of the trunk. Diaphragm displacement via video: In experiments in which we utilized the single wire medullary electrode (4/17) the data acquisition system was very sensitive to noise, and so we utilized a non-contact measure of respiration via displacement of the abdominothoracic region via video. A dark, distinguishable mark was made on the animal’s right caudal thorax such that it contrasted with the animal’s fur (Long-Evans are pied). In post-processing with ImageJ, the region of the mark was selected within the frame. A plot profile of intensity change was made through the stack of video TIFF files. Each time the animal’s diaphragm moved with respiration, the ROI encompassing the mark moved and the resulting displacement was calculated. This intensity change was registered as an analogue to diaphragmatic deflection. This technique was more effective at measuring large deflections associated with gasps and sighs and less effective with rapid shallow breathing typical in ictal eupnea.

### 2.4 Filtering and data processing

Cardiorespiratory: Non-medullary signals were filtered and amplified using custom hardware based on the Grass p511 schematic (see Appendix for exact filter values). These signals were acquired with a National Instruments DAQ (either 6343 or 6363) at 10 kHz (well above Nyquist). We used the ECG channel to find each heartbeat using the MATLAB `findpeaks()` function. We then used these times to average and subtract the ECG artifact from other channels in a 50-beat moving window. We additionally extracted breath times from the diaphragm EMG channel and swallow times from the pharyngeal EMG channel by first acquiring the envelope with the magnitude of the Hilbert transform and then smoothing with a 125 ms Hanning smoothing window. We similarly used the `findpeaks()` function.

Neuropixels: We use Bill Karsh’s SpikeGLX and CatGT for acquisition, sync, ADC MUX correction, and concatenation (Jun et al. 2017). We used SpikeInterface (Buccino et al. 2020) for filtering and its greater control of motion estimation and correction. We attempted a variety of combinations of peak detection, localization, and motion inference, and were mostly disappointed by the results. The available motion correction algorithms are deficient in periods where there is synchronous activity across much of the probe, such as gasps. The algorithms, by default, filter out synchronous activity of nearby regions if the activity spans too great an anatomical distance, assuming that such activity must be artifact. However gasps and swallows can both produce broad recruitment of the recording tract which we believe is not entirely due to motion (see Figure 4). These broad recruitments are slightly wider than spikes (likely representing synchronous activity across many neurons in nearby tissue) but are sufficiently close in spectrum and time that they impact spike detection. In Figure 4 we do not report spikes and instead report electrographic activity without median subtraction. However in all other analyses we perform median subtraction and focus on analyses of spike rates over larger timescales – i.e., the entire 60 seconds of nasal irrigation, apnea, gasping, and return to eupnea, rather than per-gasp or per-breath analyses, which we do not find to be consistent or reliable with the currently available algorithms.

The algorithms also struggle with seizure onset. The algorithms are surprisingly good at recognizing the same neuron even when it changes its firing rate due to seizure. However during seizure new neurons become active and some previously-active neurons become inactive, and the algorithms mistake these changes in activity as motion, resulting in clear splitting errors. In most animals we use the locally exclusive peak detection, monopolar triangulation peak localization, iterative template motion estimation, and inverse distance weighting motion interpolation methods, however if these methods produced a clear error we would instead try the nonrigid motion estimation, which sometimes worked. We must manually edit the motion estimation files to exclude the death of the animal, as the rapid silencing of activity produces unreasonably large motion estimates. We then feed the new, reinterpolated file in to Kilosort 2.5 with no motion correction and remove duplicates. These steps produce an estimate of locations and times of each neuronal spike in the entire experiment, which is relatively successful. Our motion plots (and their suboptimal corrections) suggest motion of no more than 100 *μ*m over the course of each experiment, which is corrected to (by our qualitative assessment of drift maps) within 60 *μ*m of the “correct” location. Kilosort also attempts to place each neuronal spike into a cluster, ideally represented a well isolated single neuron which can greatly strengthen the analyses. These results were quite poor (likely due to poor motion correction and seizure) and not a focus of this paper. Instead, we use Phy2 to analyze the results, only discarding spikes that are clearly artifacts or duplicates. In a typical curation we kept approximately 97% of non-duplicate spikes output from Kilosort (we implemented an optional Kilosort step that already removed nearly all duplicates). We next average spikes into anatomical windows of 120 *μ*m with 50% overlap in order to address suboptimal clustering outputs due to motion estimation errors. Therefore each window might represent a handful of neurons and each spike is counted in exactly two bins. Spike rates were normalized to compensate. These windows still represent high frequency single spike activity over a small anatomical space, similar to multi unit activity via a tetrode. Even aggregated, the data is much more precise than a local field potential, which represents lower frequency population activity from a much larger tissue volume. This precision is achieved because only nearby neurons produce voltage deflections of sufficient sharpness to qualify as spikes. In some data presented we average spike rate across a larger time window to demonstrate slow changes, like seizure, but all data presented still originate from single spikes.

Medullary single wire: We used a custom ASIC amplifier (Arafat et al. 2019) acquired at 10 kHz. We additionally used a Grass AM7 audio monitor to listen to electrode activity during the experiment. These wires were used instead of Neuropixel probes in 4/17 animals and are discussed explicitly. Data should be assumed to be from Neuropixel probes unless otherwise stated.

### 2.5 Histology

We successfully obtained histological samples from 9/13 animals with Neuropixels. For histological prep, prior to insertion, we used DiI (Thermofisher #V22885), a lipophilic dye (1.5 mg/mL in isopropyl alcohol) layered onto the probe (Neuropixel, IMEC) to allow for reconstruction of the probe track. Upon explantation, we perfused the animal with 200 mL of phosphate-buffered saline and then 200 mL of 4% paraformaldehyde. We extracted the brain following fixation and placed it in a vial of fixative (4% PFA) for 24 hours, and subsequently in a vial of cryoprotectant. Fixed tissue was cleared using a cryostat and stained with DAPI for better visualization of the insertion track. Mounted slices were imaged using a Zeiss LSM 510 inverted confocal microscope. Typical histological prep suggests that perfusion should begin while the heart is still beating or extremely shortly afterward. However we are studying sudden death and wish to observe complete silencing of the brain and physiological measures. In some animals we prioritized complete silencing of physiological data and therefore could only complete a partial or no perfusion and simply removed the brain into PFA (perfusion attempt began 10 minutes or more after final breath). In other animals we ended data recording as soon as it was obvious that the trial was fatal in order to obtain fully cleared tissue (perfusion attempt began within 3 minutes after final breath). Comparing these samples suggested there was little difference in our ability to localize the probe. Therefore our sample sizes of histological samples, total data, and data available during death will be slightly different.

### 2.6 Statistics

Data are reported as mean *±* standard deviation unless otherwise noted. All statistical tests are either rank sum or signed rank, as appropriate for the data type. For single comparisons we use alpha = 0.05, and for plots with multiple comparisons (such as the bar graphs in Figure 1) we use alpha = 0.001 to accommodate. Our primary measure is spike rate, and we use the terms excite and inhibit to refer to an increase or decrease, respectively, in spike rate. We do not have any measure of neurotransmitter type in any data, and cannot comment on if firing rate changes arise from neurotransmitter excitation, inhibition, or disinhibition.

**Figure 1.**
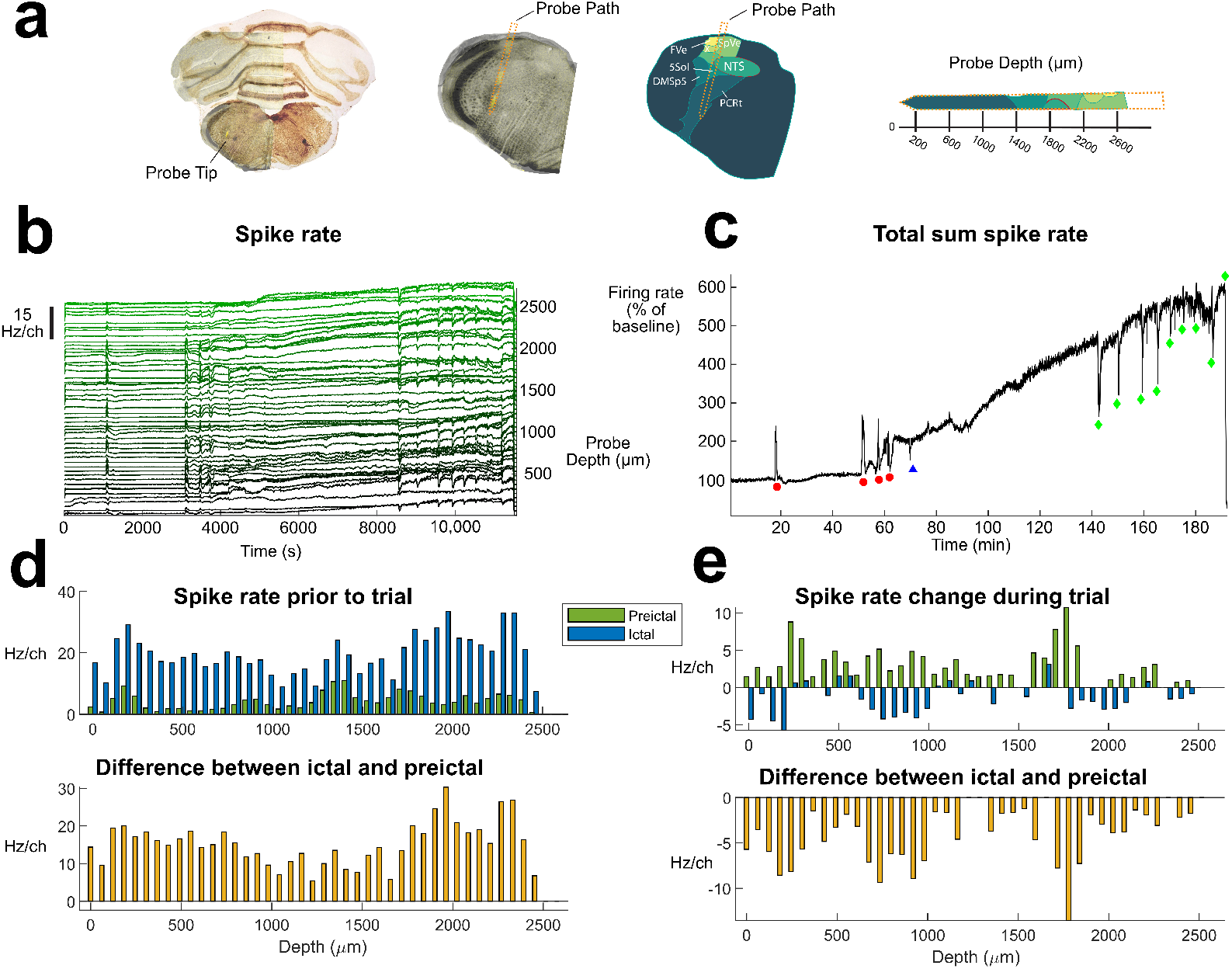
Total medullary activity at baseline, seizure, and in response to the diving reflex stimulus. a) the anatomical path of this insertion. b) and c) spike rates over the entire experiment. b) by anatomical bins and c) summed over the entire tract. Nonmedullary channels are excluded. In b) each bin spans 120 *μ*m and 12 channels, and the spike rate is normalized to the number of channels included. c) is normalized to the baseline firing rate of the first 5 minutes of data collection. Red circles indicate preictal diving reflex trials. Blue triangle indicates the time of kainic acid injection. Green diamonds indicate ictal diving reflex trials. These data indicate widespread increases in firing rate during seizure, broad excitation during preictal diving reflex trials, and broad inhibition during ictal diving reflex trials. In d) and e) we quantify firing rate during critical periods of each bin identified in b). d) the mean baseline firing rate of each channel prior to the diving reflex stimulus (n=3 preictal, n=8 nonfatal ictal), and the difference between ictal and preictal rates for each channel. The data show seizure broadly increases firing rate across the entire tissue volume. All bars shown are statistically significant p<0.001. e) the mean change in firing rate in each channel caused by the diving reflex stimulus (n=3 preictal, n=8 nonfatal ictal), and the difference in this measure between the ictal and preictal states. These data show that the post-diving-reflex-stimulus period has significantly reduced activity in the ictal state. All bar shown are statistically significant p<0.001.

## 3 Results

### 3.1 Spike rates and medullary seizure

The total activity captured by the Neuropixel probes varied dramatically from animal to animal, likely due to the variability in anatomical placement. The total spike rate across the entire region was 959 *±* 398 spikes/second, range 278–1471, n = 11 animals. For all future statements we normalize the spike rate based on the baseline period at the beginning of the experiment, calling that spike rate 100%. The injection of kainiate caused seizure onset in the medulla at approximately the same time as we would expect in cortex from prior work (Nakase et al. 2016; Sakamoto et al. 2008). In 9 / 11 animals we see a statistically significant increase in medullary firing rate in less than 15 minutes. In 2/11 animals it took up to 34 minutes to see such an increase. We tried and failed to directly record cortical onset in these experiments, as discussed in our Limitations section, so these comparisons are approximate. In most experiments medullary activity continued to increase throughout the experiment, and so the activity just before the fatal trial was 328% *±* 172%, range 146%–656%, n=11 animals, compared to baseline. There was no statistically significant correlation between the increase in firing rate and how many reflex trials the animal survived (although this comparison is imperfect as animals experienced stimuli of differing durations). When we consider the change in firing rate by anatomical location there are no obvious focal differences – almost every active region of the probe increases significantly in firing rate. An exemplary animal is shown in Figure 1. Medullary seizure does not appear to be, by itself, dangerous. Six of 11 animals experienced at least 200% activity for at least 20 minutes, including surviving multiple apnea trials. None of the animals died in seizure without an apnea trial, and no such animals were observed in the 29 animals not included in the primary dataset. Therefore animals can endure a seizure burden in the medulla for an extended period of time without danger.

### 3.2 The reflexive apnea response

Reflexive apnea trials changed the total activity in the medullary tract. For each trial we calculated the pre-trial baseline activity (to account for the higher baseline activity in seizure) and then in the subsequent 60 seconds we calculate the mean. Thus for each trial we can determine the average change induced by the reflex.

The preictal trials resulted in a 16% *±* 26% increase, (range = 17% decrease – 105% increase, n = 37 trials) in the mean activity. Conversely, ictal trials resulted in a 9.8% *±* 25% decrease (range = 71% decrease – 47% increase, n = 37 trials). These data show that ictal reflexive apnea trials are significantly more inhibitory (i.e., causing a decrease in activity) than nonseizing trials, even when we do not control for each animal and probe location (p=4e–6, rank sum, two-tailed). These results hold even when trials are paired within the same animals. In paired data we see a 33% *±* 20% decrease in activity in ictal trials (range = 70% decrease – 14% decrease, n=9 animals), which is significant (p = 0.00390, sign rank, two-tailed).

That is to say, preictal trials generally cause a moderate increase in activity in the medullary tract, although in some animals they cause a small decrease. Conversely, ictal trials generally cause a moderate decrease in activity, although in some animals they cause a small increase. When we link these comparisons per animal, then the ictal trials are always less activity than the preictal trials. Even if the ictal trial causes a small increase in activity compared to baseline, that change is always less than the increase seen preictally. An exemplary animal is shown in Figure 1. The above statements all concern the sum activity across the entire probe, but we still observe a significant change in the apnea response across the anatomical bins, as shown in Figure 1. In the 2 cases in which an ictal reflex response was observed to be excitatory the fatal trial was immediately inhibitory, not excitatory. This change is perhaps reflective of different time courses – the peak change in firing rate following a reflex trial is approximately 45 seconds after stimulus, while (discussed next) the onset of medullary shutdown is as fast as 19 seconds.

### 3.3 The central apnea response

Central apneas have been studied in this model previously, occur in a subset of animals, and are not dangerous (Nakase et al. 2016; R. B. Budde, Arafat, et al. 2018). In order to compare the impacts of central ictal apnea and reflexive ictal apnea we wanted to observe at least 3 of each apnea type of at least 3 seconds duration in a single animal. We observed such an overlap in apneas in just 1/11 animals, whose data are shown in Figure 2. These data show that, in this animal, central apneas do not cause any detectable change in the medullary activity while the reflexive apneas are strongly inhibitory. In 4 other animals we observed more examples of central apneas of shorter duration, or central apneas in which the animal died on the first or second ictal reflexive apnea, and those data all support the same conclusion.

**Figure 2.**
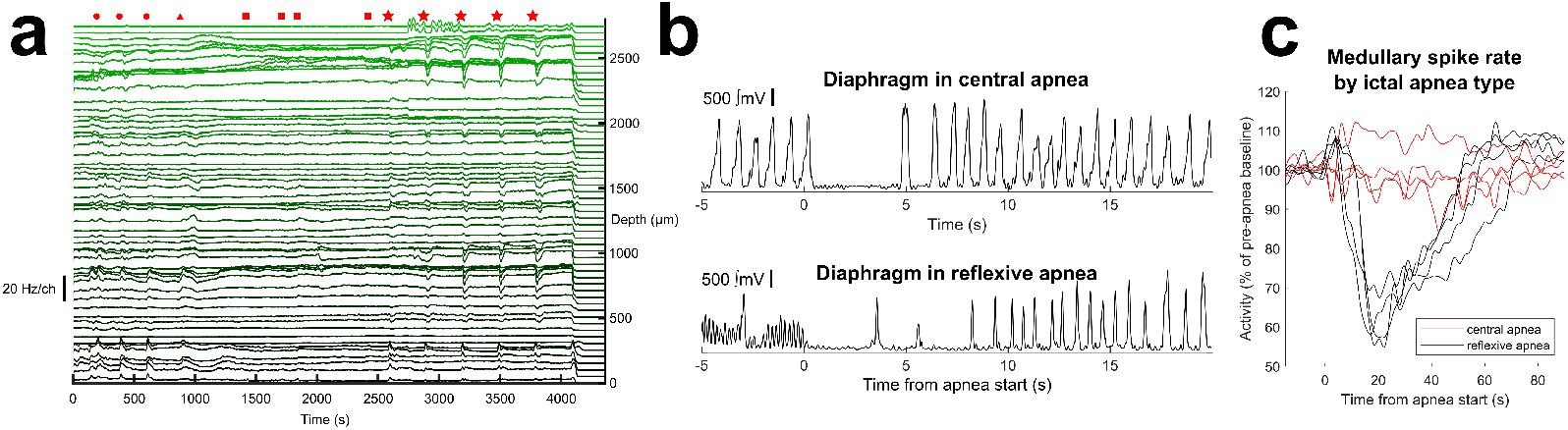
Relative impacts of central or reflexive apneas in seizure. a) outlines whole probe spike rate in anatomical bins over the entire experiment. Red circles indicated preictal diving reflex trials. Red triangle indicates the injection of kainate. Red squares indicate unprovoked central apneas of at least 3 seconds duration. Red stars indicate ictal diving reflex trials. The first such trial (first red star) did not provoke an apnea of at least 3 seconds and is not included in subfigure c) data. Across the entire probe reflexive apnea changes are obvious while central apnea changes, if present at all, are much less significant. b) example traces of integrated diaphragm activity for each apnea type. Central apneas tend to occur earlier in epileptogenesis and have a lower baseline respiratory rate which is not significantly modified before and after apnea. We provoke reflexive apneas later in seizure during which the respiratory rate is very high and slows considerably post apnea, and then gradually increases again. c) the whole probe firing during each apnea type. Reflexive apneas produce profound changes while central apneas do not (p<0.05).

### 3.4 Medullary shutdown in death

In our prior work in this model we observed cortical death 67.4 ± 8.5 seconds after stimulus (Biggs, R. Budde, et al. 2021). However, we can often visually tell that a trial is fatal much sooner, anecdotally around 40 seconds or so. During fatal trials we observe a marked increase in tonus across the animal, which is easily observable by recording EMG from any noncardiorespiratory muscle. In this experimental set we observe rapid shutdown across the entire medullary tract in fatal trials. Figure 3 shows examples of various data windows during nonfatal and fatal trials. When we consider the approximate anatomical regions of these data windows we observe no obvious trends in a single animal or across animals – the entire medullary tract appears to shut down within a narrow time range. If we consider the entire medullary tract in total then we observe death (here we define as a 90% decline in total spikes detected) within 46.7 *±* 8.4 seconds (range 34.7–59.2, n=7). However we also see some data windows shut down more rapidly (See Figure 3c). We can use these data to predict if a trial will be fatal a little faster. First, we classify candidate data channels as having firing rate more than 1 spike/second/channel and never have a firing rate of 0 spikes/second across the entire experiment (prior to death). We then determine the earliest time that one of these candidate channels falls to 0 spikes/second and remains at 0 spikes/second for the remainder of data collected. Using these “predictor” data windows we can determine that a trial is fatal 33.1 *±* 9.8 seconds (range 19.0–50.2, n=11) after the stimulus begins. The mismatched sample numbers are due to our length of data recording after death in order to obtain histological samples. In 4/11 animals we recorded sufficient data to observe multiple individual channels decline to 0 firing rate for the predictor data metric, but not sufficient data to observe decline across enough of the probe to accurately calculate the total activity metric. Note that, especially preictally, it is quite common for the apnea duration to be longer than the 33 seconds time-to-death observed in select medullary traces. These data suggest the mechanism of death cannot be suffocation alone and instead is driven by an active method of inhibition.

**Figure 3.**
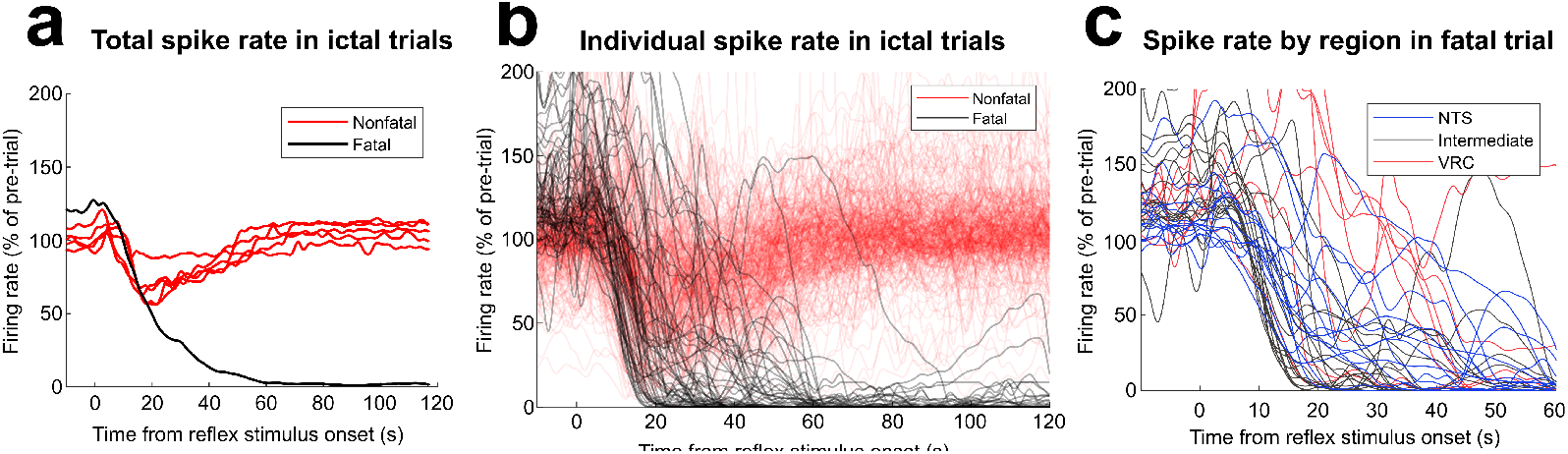
Spike rates in fatal trials. All data in this figure come from a single animal, the same animal as Figure 2. a) The spike rate across the entire probe in all ictal diving reflex trials. The fatal trial shows clear separation from nonfatal trials beginning at approximately 20 seconds after stimulus. b) the spike rate for each bin across every ictal diving reflex trial (see Figure 2a for bin rates for the entire experiment). The trends of the entire probe are well reflected in the more anatomically discrete data. c) The spike rate by anatomical bin for the fatal trial only, with labeling based on the approximate anatomical region. There are no statistically significant differences in time-to-decline based on anatomical region.

### 3.5 Swallowing and population bursts

#### 3.5.1 Single medullary wire

We performed experiments in 4 animals with a single medullary electrode. Our recording system for this electrode was more sensitive to noise, and so we primarily used video to assess diaphragmatic activity in these animals. While these data might appear limited compared to Neuropixels experiments, the 125 *μ*m silver wire offers a few properties that address some weaknesses of the Neuropixel electrodes. The larger electrode surface area presents with greater capacitance and will integrate signals over a much larger tissue volume. This integration is distinct from the averaging we use of the smaller 12 *μ*m Neuropixel electrode sites. Neuropixel sites well sample our tissue tract in the dorsoventral axis but not in the mediolateral or anteroposterior axes. Therefore the wire captures a larger tissue volume of the solitary nucleus and provides a better estimate of the synchronous firing of a larger population of neurons, while sacrificing some anatomical precision and sampling outside the NTS. Further small electrodes sites are much more sensitive to motion issues, and so the larger electrodes help support some of the phenomena we observe are distinct from artifact.

In all 4 experiments we found that the NTS presents with stable, tonic activity in eupnea, and produces no noticeable bursts with breathing. During the water stimulus and proceeding apnea there is also no significant change. However, when respiration restarts with gasping there are large population bursts in time with each gasp. These bursts gradually decline in amplitude as the gasping transitions back to eupnea. However during seizure this association changes dramatically. During an ictal water stimulus, which produces apnea and during which the NTS was previously silent, the NTS population now bursts frequently, usually accompanied by swallows. Interestingly, the bursts then are *absent* in subsequent gasping. Therefore this region shifts association from gasping towards swallowing. These observations might be impacted by motion artifact, however the fact that preictally there are not swallows and there are population bursts with gasps, and then ictally the bursts are associated with swallows and not gasps imply that motion artifact is not the sole explanation.

We were able to reproduce these results in 2 experiments with Neuropixel probes. While increased swallowing was obvious in 11/13 Neuropixel experiments, in only 2/11 did we observe the population bursts which again demonstrated the change in association like with the single 125 um wires. In these animals we demonstrate 1) clear anatomical bands of activity, which further suggest the result is not motion artifact (as such artifact would be observable across the entire probe) and 2) validation of swallowing via direct laryngeal complex recordings. These data are shown in Figure 4. We additionally observed 2/13 Neuropixel experiments in which we confirmed a **lack** of swallowing behaviors despite apneas and death. Therefore these observations represent a phenotype which is a subset of all cases and is not necessary for death. However these data implicate the control of swallowing in the network architecture of reflexive apneas and their abnormal control in seizure.

**Figure 4.**
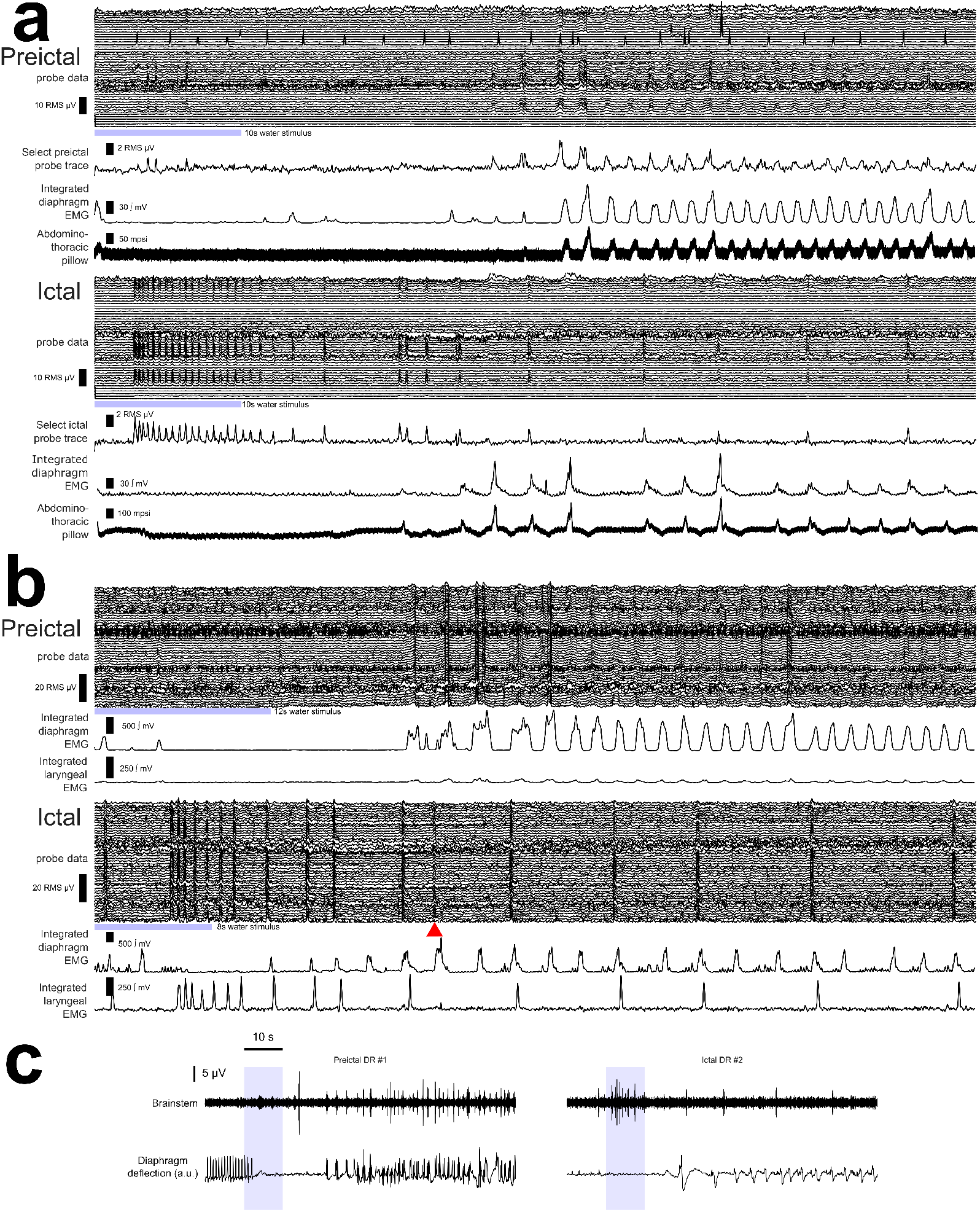
Population bursts and swallowing follow reflexive apnea. This figure demonstrates both a phenotypic phenomenon (swallowing and population bursts) as well as shortcomings of the available data analysis methods. The probe data in this figure undergo filtering but not motion correction or median subtraction, which both destroy the phenomenon in question (or migrate it to a different region of tissue, which is equally problematic). Each subset, a, b, c, represent data from a single, unique animal. In a) and b) the probe data show binned (120 *μ*m) smoothed (50 ms) electrographic activity, with blue bars showing water stimulus. All other measures as described in the methods. a) Data from a single animal from representative preictal and ictal apnea trials. In the preictal case probe data show medullary population bursts with gasps following the apnea. Population bursts slightly lead diaphragm displacement and are well correlated. Ictally, this correlation is lost, with population bursts only occasionally overlapping gasps. In all cases there is a clear anatomical band of activity on the probe which is the same in preictal and ictal cases. This separation of activity, as well as the change in correlation between gasps and bursts, strongly suggest this activity is not due to motion artifact. b) A second example, similar to a), but in a case in which swallow activity is recorded. This animal displays less robust anatomical band separation, but still shows probe population bursts are correlated with gasps preictally and correlated with swallows ictally. The red triangle denotes the single ictal exception to this observation. Note that in this animal swallows follow after gasps when they are paired. This observation is not true in all animals, but is consistent within the same animal (i.e., if swallows proceed gasps then they always do in that animal, while in others they may precede). c) This animal demonstrates the same phenomenon as a) and b) but in an animal with the single 125 *μ*m wire. This wire records from a much larger population of neurons than the Neuropixel sites. These data reinforce the observations of a) and b). In both a) and b) these population bursts will be removed if using standard Neuropixel median subtraction and motion correction tools, as these assume that neurons across the probe are mostly uncorrelated. Indeed, in animal a), median subtraction results in population bursts being shifted to the quiet anatomical bands, creating artifactual phantom activity in a region that is actually silent throughout the experiment (and this activity is then identified as spikes and clustered as a high quality single unit neuron).

## 4 Discussion

These results suggest:

1. Seizure (in this chemoconvulsant status model) spreads to the medulla at approximately the same rate and intensity as cortex shown in our prior work. However it is also safe, as animals can endure high levels of activity for hours without mortality. Medullary seizure is only fatal when paired with a reflexive apnea stimulus.
2. The reflexive apneas produce a more inhibitory response in the medullary tract in seizure compared to baseline. While our data is limited, it suggests that central apneas produce no detectable response. Therefore reflexive apnea may represent a unique apnea type that interferes more powerfully in medullary signaling, and may be more destabilizing.
3. The entire medullary tract shuts down quite rapidly in every fatal apnea. Some “predictor” channels shut down in less than 20 seconds after the stimulus starts. These data suggest the death is the result of active inhibition pathways. Suffocation alone cannot explain such a rapid, widespread decline in activity. These data are perhaps supportive of the theory of a spreading wave of depolarization (Aiba and Noebels 2015).
4. Some data suggest swallowing emerges as an ictal phenotype which competes for gasping drive. The counteractive roles of swallowing (anti-inspiratory) and gasping (pro-inspiratory) may play a role in sudden respiratory deaths more broadly.

Our data demonstrates that seizure spread to the medulla might be necessary but is not sufficient for ictal apneic deaths. Rather, we require two factors – a network impairment (seizure in this study, or as we showed previously the unilateral or bilateral resection of the carotid bodies) and the activation of a strong apneic stimulus (in this work the nasal irrigation, but previously also laryngeal pH chemosensation, and also possibly specific forebrain-initiated apneic responses).

Predictions:

1. The activation of a reflexive apnea during seizure may cause a wave of spreading depolarization as described in (Aiba and Noebels 2015), or may lower the threshold for a subsequent spreading depolarization event.
2. Most ictal central apneas in generalized seizure should not provoke spreading depolarization events.
3. Forebrain seizure that induces dangerous apneas (near-SUDEP, of extended duration, and/or producing cyanosis) may initiate overlapping apnea pathways as the reflexive apneas of this work.
4. The significant literature of diving reflex and apnea plasticity imply that long-lasting, trained changes in this network may shift the threshold for sudden death.

## 5 Limitations

Population bursts and swallowing: We only observed populations bursts correlated to gasps and/or swallows in a subset of animals. In each case we observed the transition of bursts from gasp associated preictally to swallow associated ictally. In some animals we observed no swallows at all. Swallowing is inhibitory to respiration and may contribute to the fatal decline in respiratory drive, but swallowing as a behavior is not necessary for death. Rather the emergence of swallowing may represent an outward sign of changes in the medullary signaling balance between different respiratory behaviors. When swallows occur we do not always record population bursts, which may be a limitation of our anatomical animal-to-animal variability in recording locations. Future work with more precise histological methods may produce more consistent results.

Speed of medullary decline vs cortical decline: We mention that our data are suggestive that the medullary centers silence before the cortex, but these data are imperfect. In these experiments we were unsuccessful at recording electrocorticography simultaneous to the brainstem electrodes. Our ECoG recording setup infected the Neuropixel medullary data with so much noise that the Neuropixel data was unusable. Therefore we must rely on data from a prior experimental set to make such a comparison. Further, we are comparing lower frequency population events (ECoG) to individual spike counts from a Neuropixel probe. These data are similar but not identical in type, and so direct comparisons may be inaccurate. We do not have the ability to record from more than one Neuropixel probe at once, but a more appropriate solution would be to use one cortical probe and one medullary probe and compare spike counts directly.

Central vs obstructive apneas: Central apneas in this model only occur in about 50% of animals over several hours. Due to our large dataset limits (>300 GB per animal) we do not record for many hours to observe these specifically. For a “fair” comparison between apnea types we also require central apneas to be of sufficient duration compared to the reflexive apneas (we have more examples of central apneas <3 seconds in duration which do not cause inhibition, but these are perhaps not comparable to reflexive apneas of >8 seconds duration). Further, for sufficient comparisons we require multiple examples (>3 per animal) of both apnea types, and seizing animals will often die too quickly of reflexive apneas. The intersection of these limitations results in strong comparisons between apnea types being quite rare (only succeeding in 1/11 attempts). A more thorough comparison of these apneas will require greater data capabilities (>1 TB per animal) and would be aided by a method of resuscitation from fatal reflexive apnea to ensure adequate per-animal sample sizes.

## A Appendix

### A.1 Filtering

**Table 1.**
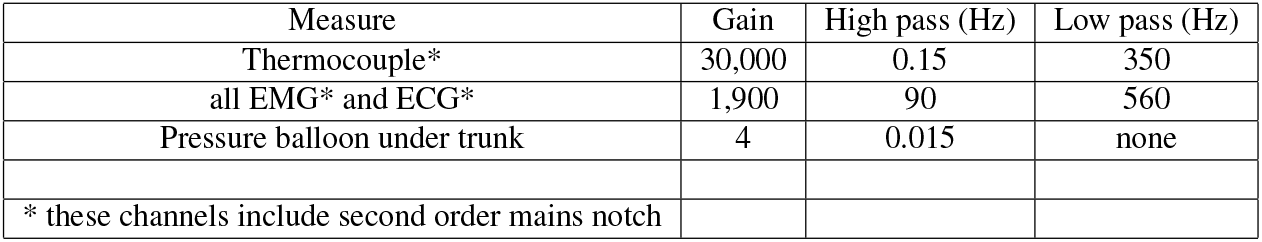
Filter settings.

| Measure | Gain | High pass (Hz) | Low pass (Hz) |
| --- | --- | --- | --- |
| Thermocouple* | 30,000 | 0.15 | 350 |
| all EMG* and ECG* | 1,900 | 90 | 560 |
| Pressure balloon under trunk | 4 | 0.015 | none |
| * these channels include second order mains notch |  |  |  |

## Author contributions statement

**Ryan Budde**: Conceptualization of this study, Methodology, Software, Analysis, Writing. **Laura RoaFiore**: Conceptualization of this study, Methodology, Surgery, Analysis, Writing. **Pedro Irazoqui**: Conceptualization of this study, Coordinator, Review.

## Acknowledgments

This work was funded by NIH NS119390. The authors have no conflicts of interest to disclose. Data is available with reasonable request.

